# Genotype-phenotype correlation analysis and therapeutic development using a patient stem cell-derived disease model of Wolfram syndrome

**DOI:** 10.1101/2021.11.07.467657

**Authors:** Rie Asada Kitamura, Kristina G. Maxwell, Wenjuan Ye, Kelly Kries, Cris M Brown, Punn Augsornworawat, Yoel Hirsch, Martin M Johansson, Tzvi Weiden, Joseph Ekstein, Joshua Cohen, Justin Klee, Kent Leslie, Anton Simeonov, Mark J. Henderson, Jeffrey R. Millman, Fumihiko Urano

## Abstract

Wolfram syndrome is a rare genetic disorder largely caused by pathogenic variants in the *WFS1* gene and manifested by diabetes mellitus, optic nerve atrophy, and progressive neurodegeneration. Recent genetic and clinical findings have revealed Wolfram syndrome as a spectrum disorder. Therefore, a genotype-phenotype correlation analysis is needed for diagnosis and therapeutic development. Here, we focus on the *WFS1* c.1672C>T, p.R558C variant which is highly prevalent in the Ashkenazi-Jewish population. Clinical investigation indicates that subjects carrying the homozygous *WFS1* c.1672C>T, p.R558C variant show mild forms of Wolfram syndrome phenotypes. Expression of WFS1 p.R558C is more stable compared to the other known recessive pathogenic variants associated with Wolfram syndrome. Stem cell-derived islets (SC-islets) homozygous for *WFS1* c.1672C>T variant recapitulates genotype-related Wolfram phenotypes, which are milder than those of SC-islets with compound heterozygous *WFS1* c.1672C>T (p.R558C), c.2654C>T (p.P885L). Enhancing residual WFS1 function by a combination treatment of chemical chaperones, sodium 4-phenylbutyrate (4-PBA) and tauroursodeoxycholic acid (TUDCA), mitigates detrimental effects caused by the *WFS1* c.1672C>T, p.R558C variant and restored SC-islet function. Thus, the *WFS1* c.1672C>T, p.R558C variant causes a mild form of Wolfram syndrome phenotypes, which can be remitted with a combination treatment of chemical chaperones. We demonstrate that our patient stem cell-derived disease model provides a valuable platform for further genotype-phenotype analysis and therapeutic development for Wolfram syndrome.

**One sentence summary:** Development of personalized therapy for Wolfram syndrome using genetics and iPSC model.

## Introduction

Wolfram syndrome is a rare, monogenic life-threatening disease largely caused by pathogenic variants in the Wolfram syndrome (*WFS1*) gene or, in a small fraction of patients, pathogenic variants in the CDGSH iron sulfur domain protein 2 (*CISD2*) gene (1–3). There is currently no treatment to delay, halt, or reverse the progression of this disease. Wolfram syndrome is well characterized by juvenile-onset insulin-dependent diabetes, optic nerve atrophy, and progressive neurodegeneration (4, 5). Many patients also develop other symptoms, ranging from hearing loss and endocrine deficiencies to neurological and psychiatric conditions (4, 6). Accordingly, recent clinical and genetic findings have revealed that Wolfram syndrome is best characterized as a spectrum disorder (7). Of the approximately 200 *WFS1* variants associated with Wolfram syndrome, approximately 35% are missense, 25% are nonsense, 21% are frameshift, 13% are in-frame insertion or deletions, and 3% are splice-site variants (8, 9). Most of these variants are predicted to be inactivating, loss-of-function variants, but extensive molecular characterization of individual alleles is sparse. Hence, there is a great need for genotype-phenotype correlation data to guide diagnostic interpretation of *WFS1* variants.

*WFS1* encodes an endoplasmic reticulum (ER) transmembrane protein. The ER is a central cell organelle responsible for protein folding, Ca^2+^ storage, and lipid synthesis. It has been reported that WFS1 regulates Ca^2+^ homeostasis in the ER, which is crucial in the synthesis and secretion of neurotransmitters and hormones such as insulin (10, 11). WFS1 deficiency in the ER causes Ca^2+^ homeostasis disruption, leading to chronic ER stress followed by the unfolded protein response (UPR) (12, 13). WFS1 also negatively regulates ATF6, a UPR molecule, inhibiting hyperactivation of ATF6 and consequent cell apoptosis (14). Furthermore, a recent study suggested that WFS1 impacts mitochondrial function by transporting Ca^2+^ from the ER to the mitochondria via the mitochondria-associated ER membrane (MAM) (15).

Several *Wfs1* knockout rodents were developed as disease models of Wolfram syndrome, which generate insight into etiology and provide opportunities to test therapeutic agents (11, 16–19). The models display progressive glucose intolerance due to impaired glucose-stimulated insulin secretion (GSIS) and increased pancreatic *β* cell death (16, 18–20). However, the onset of diabetes in these rodent models is delayed relative to the human phenotype, with further variation between each model based on rodent strain (16, 19). Also, to the best of our knowledge, there are no transgenic animals with phenotypes and variants corresponding to the pathogenic *WFS1* variants found in patients with Wolfram syndrome. As a result, these rodent models may not fully capture the spectrum of Wolfram phenotypes. By contrast, patient induced pluripotent stem cells (iPSCs) differentiated into disease-relevant cell types have been demonstrated as suitable models for genotype-phenotype correlation analysis (21, 22).

To date, significant efforts have been invested to develop novel Wolfram syndrome treatments (23). Compounds such as valproic acid and glucagon-like peptide (GLP)-1 receptor agonists were identified as possible drug candidates based on preclinical studies in immortalized cell and rodent models (24–26). In addition, we recently conducted a phase 1b/2a clinical trial of dantrolene sodium, an ER Ca^2+^ stabilizer, which demonstrated efficacy in a subset of patients with Wolfram syndrome (11, 27). However, identifying additional therapeutic candidates for patients with Wolfram syndrome is still needed.

Here, we focus on the missense variant, *WFS1* c.1672C>T, p.R558C, which is enriched in the Ashkenazi Jewish population (allele frequency 1.4%) (28). We characterize this variant multidimensionally through clinical investigation, biochemical studies, and patient iPSC-derived disease models. Further, we demonstrate the potential efficacy of a combination treatment of chemical chaperones, sodium 4-phenylbutyrate (4-PBA) and tauroursodeoxycholic acid (TUDCA), as a novel therapeutic approach for Wolfram syndrome.

## Results

### *WFS1* c.1672C>T, p.R558C is enriched in Ashkenazi Jewish population and causes a mild form of Wolfram phenotypes

To determine the carrier frequency for *WFS1* c.1672C>T, p.R558C variant in the Jewish population, we genotyped 87,093 subjects from several Jewish populations. In the original data set, each subject was classified by self-identification as Ashkenazi, Sephardi, Ashkenazi/Sephardi, Convert, and Unknown. Samples from Converts and Unknown origin made up a total of 773 and were excluded from analysis. The observed frequency of *WFS1* c.1672C>T, p.R558C carriers in Ashkenazi Jewish subjects reached 2.32% (1:43), 1.32% (1:76) for the Ashkenazi/Sephardi, and 0.04% (1:2,268) in Sephardi Jewish subjects (Figure 1A and Supplemental Table S1A). To elucidate if *WFS1* c.1672C>T, p.R558C was present at higher rates in the Jewish population residing from various countries, we classified data based on self-reported ancestry of four grandparents. Subjects who stated two or more countries of mixed origin were removed from analysis. In cases where South Africa was provided as the country of origin, the samples were redefined as Lithuanian as South African Jews are primarily of Lithuanian origin (29). Subjects who had Israel or USA stated in their ancestry were also removed because the Jewish people residing in these countries often have mixed Ashkenazi origins (30). Subjects that did not provide any information on grandparental origin or stated Unknown were removed. Subjects with Ukrainian origin were merged into the Russian group. Subjects with Belarus and Czechia origin were removed from analysis because they totaled less than 100 subjects and a few carriers in small sample group tend to produce spurious signal. In data classified by the country of origin, the frequencies occur as follows: Romania 3.50% (1:29), Poland 2.57% (1:39), Russia 2.07% (1:48), Hungary 1.63% (1:61), Germany 1.60% (1:63) and Lithuania 0.87% (1:116) (Figure 1B and Supplemental Table S1B). Clinical investigation revealed that most subjects carrying the homozygous *WFS1* c.1672C>T, p.R558C variant developed diabetes mellitus, however, the age at diagnosis was greater than that of typical Wolfram syndrome (approximately 6 years) (4) (Figure 1C and D). Only four subjects were clinically diagnosed with optic nerve atrophy. Their optic nerve atrophy is mild, and no case was diagnosed legally blind (Figure 1C and D). Additionally, no subject developed hearing loss nor diabetes insipidus (Figure 1C and D). Together, *WFS1* c.1672C>T, p.R558C variant is enriched in the Ashkenazi Jewish population, especially those originated from Romania, and the variant leads to mild or less severe phenotypes of Wolfram syndrome.

**Figure 1.**
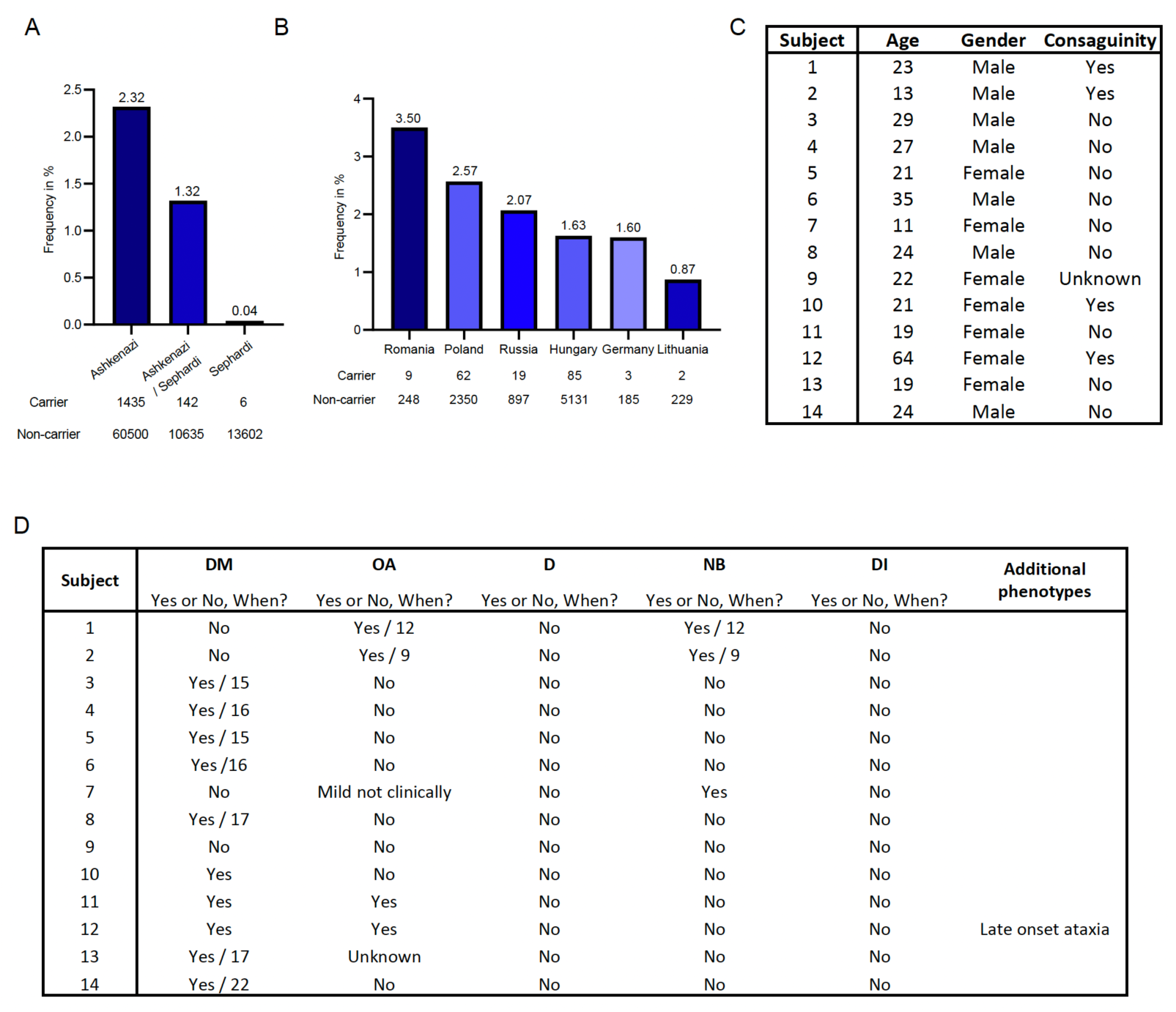
Carrier frequencies and clinical manifestation of *WFS1* c.1672C>T, p.R558C variant. (A) Carrier frequencies for *WFS1* c.1672C>T, p.R558C in subjects of Ashkenazi, Ashkenazi/Sephardi and Sephardi descent. (B) Carrier frequencies for *WFS1* c.1672C>T, p.R558C by country origin. (C) Information of subjects with homozygous *WFS1* c.1672C>T, p.R558C variant. (D) Clinical manifestation of *WFS1* c.1672C>T, p.R558C homozygotes. DM: Diabetes Mellitus, OA: Optic nerve atrophy, D: Deafness, NB: Neurogenic Bladder, DI: Diabetes Insipidus.

### The WFS1 p.R558C variant is degraded more than wild-type, but less than WFS1 p.P885L variant

Pathogenic *WFS1* variants are classified based on their effect on WFS1 expression: class A, depleted WFS1 protein or reduced, defective WFS1 protein, which leads to loss-of-function or incomplete function; class B, expression of defective WFS1 protein leading to gain-of-function. Class A is furthermore divided into three subclasses: class A1, WFS1 depletion due to *WFS1* mRNA degradation (Nonsense Mediated Decay, NMD); class A2, WFS1 depletion due to WFS1 protein degradation; class A3, WFS1 depletion due to mRNA and protein degradation (31, 32)(Supplemental Figure 1A). To determine the class specification of the *WFS1* c.1672C>T, p.R558C variant, we investigated the thermal stability of WFS1 p.R558C and p.P885L by appending a HiBiT-based tag to detect the variant in cells (33). The p.R558C variant showed less thermal stability than wild-type WFS1, suggesting an altered folding state, but more stability compared to the known autosomal recessive variant p.P885L which is pathogenic and is associated with a typical form of Wolfram syndrome (34, 35)(Figure 2, A and B). Both p.R558C and p.P885L expression could be rescued by incubating cells at reduced temperature, supporting a folding defect conferred by the variants (Figure 2C). Treatment with a proteasome inhibitor, bortezomib, increased WFS1 protein from both variants and the fold change for p.P885L was higher than p.R558C (Figure 2D), indicating that proteasomal degradation of p.R558C is less than p.P885L. To confirm this observation, we performed a cycloheximide (CHX) chase assay using HA-tagged WFS1 variants. After inhibiting translation of nascent protein by CHX treatment, the protein levels of p.R558C and p.P885L were rapidly decreased within 2 hours (Figure 2E). However, the rate of p.P885L decay was faster than p.R558C (Figure 2E). Also, the basal expression of p.P885L was lower before CHX treatment compared to WT and p.R558C, all consistent with more rapid degradation of p.P885L (Figure 2E).

**Figure 2.**
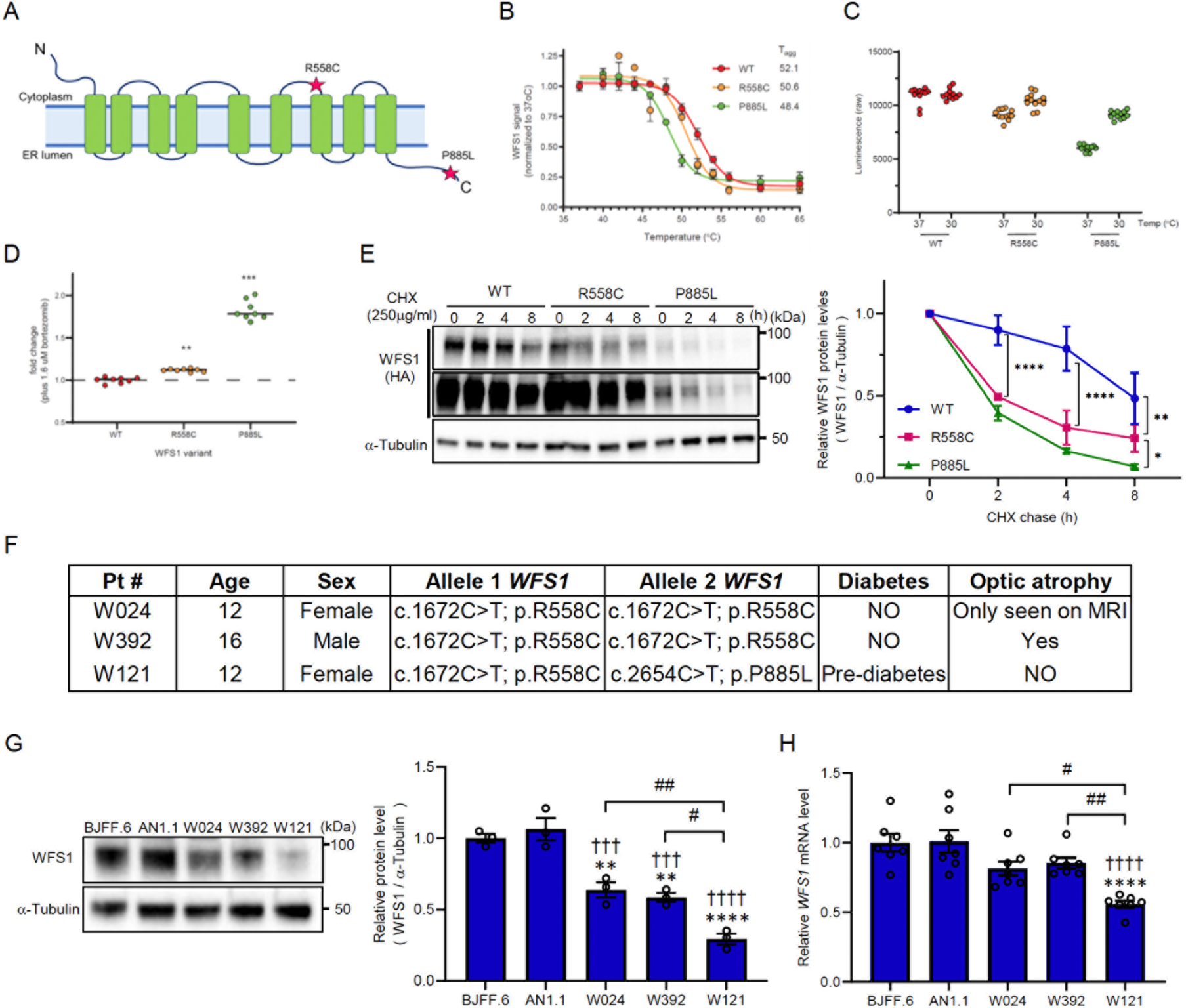
WFS1 p.R558C is more stable in the cell compared to p.P885L variant. (A) Diagram of WFS1 protein showing the location of two variants, R558C and P885L. (B) (Left) Thermal profiles of WFS1 variants (WT, R558C and P885L) measured using SplitLuc tagged reporters expressed in HEK293T cells. (C) Luminescence intensities of WFS1 variants in cells incubated at 30 and 37°C for 24 h. (D) Fold change of luminescence intensities of WFS1 variants treated with proteasome inhibitor, bortezomib, for 24 hours. (E) (Left) Representative blotting image of WFS1 (HA) and a-Tubulin in CHX chase assay. Lower panel of WFS1 (HA) is long-exposure image. (Right) A quantification of relative WFS1 protein level normalized with *α*-Tubulin. (n=3, *P<0.05, **P<0.01 and ****P<0.0001 by two-way ANOVA). (F) Information on the three patients, including the genetic location of autosomal recessive pathogenic variants in *WFS1* and symptoms. (G) (Left) Representative blotting image of WFS1 and *α*-Tubulin in iPSCs. (Right) A quantification of relative WFS1 protein level normalized with *α*-Tubulin (n=3, **P<0.01 and ****P<0.0001 by one-way ANOVA compared to BJFF.6, †††P<0.001 and ††††P<0.0001 by one-way ANOVA compared to AN1.1, #P<0.05 and ##P<0.01 by one-way ANOVA). (H). Relative mRNA level of *WFS1* in iPSCs. (n=7, ****P<0.0001 by one-way ANOVA compared to BJFF.6, ††††P<0.0001 by one-way ANOVA compared to AN1.1, #P<0.05 and ##P<0.01 by one-way ANOVA)

Next, we examined if the *WFS1* variants endogenously expressed in cells would show similar post-translational stabilities. We obtained peripheral blood mononuclear cells (PBMCs) from three subjects carrying pathogenic variants in the *WFS1* gene (W024: c.1672C>T, c.1672C>T; W392: c.1672C>T, c.1672C>T; W121: c.1672C>T, c.2654C>T) and generated iPSCs (Figure 2F and Supplemental Figure 1B). Consistent with our clinical investigation, subjects W024, W392, and W121 had mild phenotypes of Wolfram syndrome (Figure 2E). Western blot (WB) analysis revealed a reduction in WFS1 protein levels for W024, W392 and W121 compared to two control iPSC lines (BJFF.6 and AN1.1) (Figure 2F). Of the three patient lines, WFS1 protein level in W121 was less than W024 and W392 (Figure 2F). *WFS1* mRNA was not significantly decreased in W024 and W392 compared to control lines but was reduced for W121 (Figure 2G). Taken together, the *WFS1* c.1672C>T, p.R558C variant leads to reduced expression of defective WFS1 protein that is driven by post-translation protein degradation, but not mRNA alterations, designating the variant as class A2.

### A combination treatment of 4-PBA and TUDCA ameliorates cellular function in neural progenitor cells with c.1672C>T, p.R558C variant

Wolfram syndrome is recognized as an ER disorder (6, 36, 37). Given that the ER and mitochondria interact both physiologically and functionally to maintain cellular homeostasis and determine cell fate under pathophysiological condition, pathogenic *WFS1* variants cause not only ER dysfunction but also dysregulation of mitochondrial dynamics and appropriate function (38, 39). This evidence suggests that a combination drug modulating multiple targets simultaneously in the cell could be a good treatment candidate for Wolfram syndrome. Chemical chaperones, such as 4-PBA and TUDCA, are well known to rescue or stabilize the native conformation of proteins by interacting with exposed hydrophobic segments of the unfolded protein (40). In addition to its chaperone activity, 4-PBA exhibits HDAC inhibitory activity, which transcriptionally induces the expression of molecular chaperones (41). TUDCA has been reported to reduce reactive oxygen species (ROS) formation (42), prevent mitochondrial dysfunction (43), and inhibit apoptosis through the intrinsic (44) and extrinsic pathways (45). It has been shown that 4-PBA can improve insulin synthesis in Wolfram iPSC-derived *β* cells (46). We, therefore, hypothesized that a combination treatment of 4-PBA and TUDCA would have an additive effect on the restoration of reduced WFS1 expression and organelle dysfunction. Specifically, the combination treatment would be expected for effects on pathogenic *WFS1* variants that are degraded mainly at the protein level, such as WFS1 p.R558C (Figure 3A).

**Figure 3.**
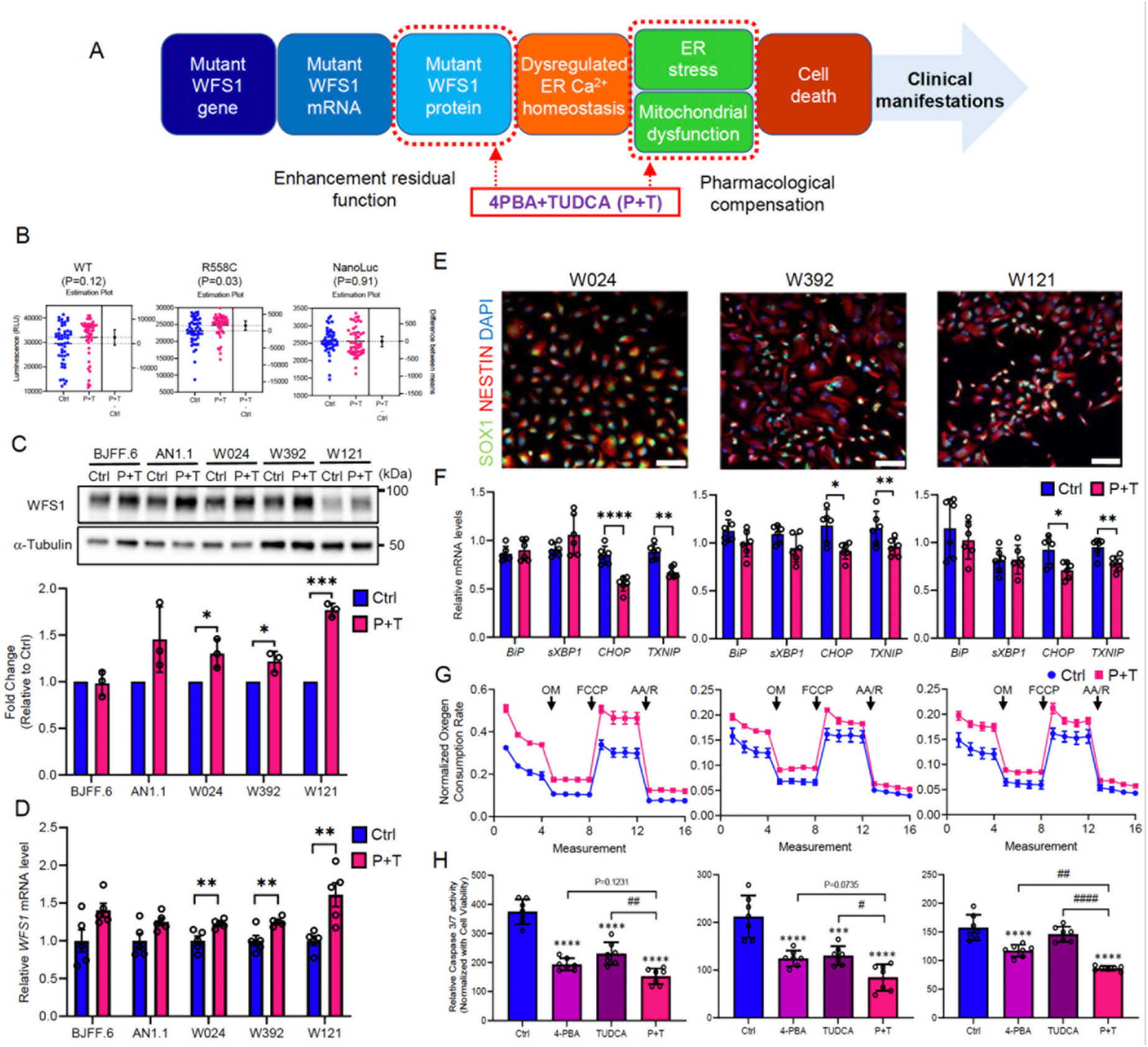
A combination treatment with 4-PBA and TUDCA mitigates detrimental effect of *WFS1* c.1672C>T, p.R558C variant. (A) A schematic of Wolfram syndrome etiology and the targets to modulate by a combination treatment of 4-PBA and TUDCA (P+T). (B) Expression of HiBiT tagged WFS1 protein after 24 h treatment with 500 μM 4-PBA and 50 μM TUDCA (P+T). NanoLuc levels, expressed from an identical plasmid backbone, were examined. (C) (Upper) Representative blotting images of WFS1 and *α*-Tubulin in iPSCs treated with or without P+T for 48 hours. (Lower) A quantification of WFS1 protein levels normalized with *α*-Tubulin. (n=3, *P<0.05 and ***P<0.001 by unpaired t-test compared to Ctrl). (D) Relative mRNA levels of *WFS1* in iPSCs treated with or without P+T for 48 hours (n=5, **P<0.01 by unpaired t-test compared to Ctrl). (E) Representative immunofluorescence images of neural progenitor cell (NPC) markers in NPCs differentiated from patient-derived iPSCs. Scale bar: 100 μm. (F) qPCR analysis of ER stress related genes in NPCs treated with or without P+T for 48 hours. (n=6, *P<0.05, **P<0.01 and ****P<0.0001 by unpaired t-test compared to Ctrl). (G) Mitochondrial respiration of NPCs treated with or without P+T for 48 hours represented as percentage of baseline oxygen consumption rate measurements. Respiration was interrogated by measuring changes in relative OCR after injection with oligomycin (OM), FCCP, and antimycin A (AA)/rotenone (R). (n=3, W024: ****P<0.0001, W392: ****P<0.0001, and W121: ****P<0.0001 by tow-way ANOVA) (H) Caspase 3/7 activity normalized with cell viability in NPCs treated with or without either of 4-PBA, TUDCA and P+T for 48 hours. (n=7, ***P<0.001 and ****P<0.0001 by one-way ANOVA compared to Ctrl, #P<0.05, ##P<0.01 and ####P<0.0001 by one-way ANOVA).

We first tested if a combination treatment with 4-PBA and TUDCA (P+T) stabilized WFS1 protein using the HiBiT tagged reporters. The incubation with P+T significantly increased the steady-state levels of WFS1 p.R558C protein, but not WT nor a NanoLuc control expressed from an identical plasmid backbone (Figure 3B). We also screened the NCATS Pharmaceutical Collection, which includes approved drugs (∼2000 compounds) and found a small number of compounds that increase WFS1 p.R558C protein level, of which disulfiram was the top hit, but magnitude of effect was similar to the P+T (Supplemental Figure 2 and Supplemental Data 1). We also compared endogenous WFS1 protein levels in iPSCs treated with P+T. The P+T treatment significantly increased WFS1 protein levels in iPSCs derived from all three patient lines (Figure 3C). Additionally, mRNA level was increased by the P+T treatment (Figure 3D). We previously described organelle dysfunction followed by cell death in neural progenitor cells (NPCs) differentiated from iPSCs derived from patients with typical Wolfram syndrome (11). Therefore, we examined if a combination treatment of 4-PBA and TUDCA would restore organelle functions in NPCs derived from the three patient iPSC lines (Figure 3E). The expression of ER stress marker genes, *BiP* and *spliced XBP1* (*sXBP1*), were not affected by the P+T treatment, whereas ER stress-induced apoptosis genes, *CHOP* and *TXNIP*, were significantly decreased in each of the three patient lines (Figure 3F). We also measured the oxygen consumption rate (OCR) of NPCs to further assess mitochondrial function. Increased OCRs were observed throughout the assay in each of the three patient lines with the P+T treatment (Figure 3G). Moreover, P+T treatment inhibited apoptosis, as indicated by caspase 3/7 activity, in each of the three patient lines (Figure 3H). Of note, the inhibition by the P+T treatment was significantly more effective compared to a single treatment with TUDCA in W024 and W392 (Figure 3H). For W121 NPCs, P+T treatment was more effective compared to a single treatment of either 4-PBA or TUDCA alone (Figure 3H). In addition, we confirmed P+T treatment reduced caspase 3/7 activity in NPCs derived from patients with typical Wolfram syndrome as well (Supplemental Figure 3A and B). Together, a combination treatment of 4-PBA and TUDCA increased WFS1 expression and inhibited apoptosis by mitigating ER stress and mitochondrial dysfunction.

### A combination treatment of 4-PBA and TUDCA improves insulin secretion and survival in SC-β cells with *WFS1* c.1672C>T, p.R558C variant

The majority of patients with Wolfram syndrome develop diabetes mellitus due to the pathogenic *WFS1* variants causing detrimental effects in pancreatic *β* cells (13, 14, 20). To evaluate the impact of the *WFS1* c.1672C>T, p.R558C variant on *β* cells, we generated stem cell–derived pancreatic islets (SC-islets) from W024 and W121 iPSCs, and AN1.1 iPSCs as a control. We previously developed a 6-stage differentiation strategy, incorporating cytoskeleton modulation, to produce SC-islets containing hormone-secreting endocrine cell types, including insulin-positive stem cell-derived *β* (SC-*β*), glucagon-positive stem cell-derived α (SC-α), and somatostatin-positive stem cell-derived δ (SC-δ) cells (47) (Supplemental Figure 4). The W024 and W121 Stage 6 SC-islets produced C-peptide+ cells co-expressing *β* cell differentiation marker (NKX6.1) and committed endocrine cell marker (CHGA). The *β* cell population was similar between W024 and control SC-islets, but reduced in the W121 line (Figure 4, A and B). WFS1 protein was expressed in SC-islets derived from all three lines, with greater expression detected in control SC-islets (Figure 4C). Of note, WFS1 protein level was significantly higher in W024 SC-islets when compared to W121 (Figure 4C). We tested the functional capacity of the SC-islets in response to high glucose (20 mM) using the glucose stimulated insulin secretion (GSIS) assay. Throughout GSIS, W024 and W121 SC-islets secreted less insulin compared to control SC-islets. W024 SC-islets were able to increase their insulin secretion in response to the glucose stimulus, whereas W121 SC-islets were not capable of a glucose-stimulated response (Figure 4D). These data suggest the *WFS1* c.1672C>T, p.R558C variant has a milder effect on *β* cell function than the *WFS1* c.2654C>T, p.P885L variant.

**Figure 4.**
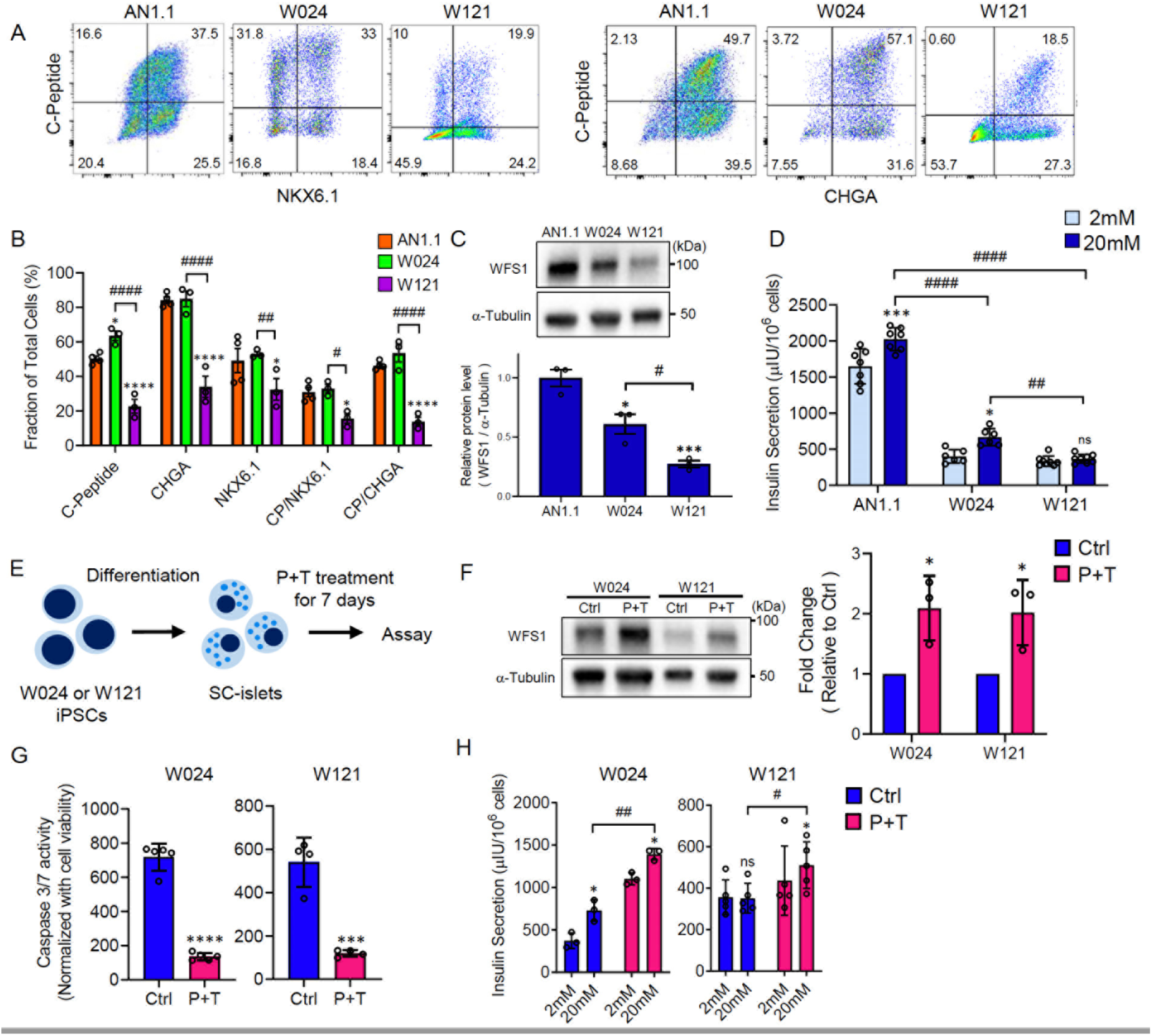
β cell function is restored by a combination treatment of 4-PBA and TUDCA in SC-islets with *WFS1* c.1672C>T, p.R558C variant. (A) Representative flow cytometry dot plots and (B) quantified fraction of cells expressing or co-expressing pancreatic *β* cell or committed endocrine cell markers for AN1.1 (n=4) W024 (n=3) and W121 (n=3) stage 6 SC-islets (*P<0.05 and ****P<0.0001 by two-way ANOVA compared to AN1.1. #P<0.05, ##P<0.01 and ####P<0.0001 by two-way ANOVA). (C) (Upper) Representative blotting image of WFS1 and a-Tubulin in stage 6 SC-islets. (Lower) A quantification of relative WFS1 protein level normalized with *α*-Tubulin (n = 3, **P<0.01 and ***P<0.001 by one-way ANOVA compared to AN1.1. #P<0.05 by one-way ANOVA). (D) Static GSIS functional assessment of AN1.1 (n = 7), W024 (n = 6) and W121 (n = 8) stage 6 SC-islets (*P < 0.05 and ***P < 0.001 by two-way ANOVA compared to 2mM of each line. ##P < 0.01 and ####P < 0.0001 by two-way ANOVA). (E) A schematic of P+T verification in SC-islets. (F) (Left) Representative blotting images of WFS1 and a- Tubulin in stage 6 SC-islets treated with or without P+T for 7 days. (Right) A quantification of WFS1 protein levels normalized with *α*-Tubulin. (n=3, *P<0.05 by unpaired t-test compared to Ctrl). (G) Caspase 3/7 activity normalized with cell viability in stage 6 SC-islets treated with or without P+T for 7 days (n=3, ***P < 0.001 and ****P < 0.0001 by unpaired t-test compared to Ctrl). (H) Static GSIS functional assessment of W024 (n = 5) and W121 (n = 4) treated with or without P+T for 7 days (*P < 0.05 by one-way paired t-test compared to 2mM of each condition. #P < 0.05 and ##P < 0.01 by two-way unpaired t-test). CP: C-peptide, CHGA: Chromogranin A, ns: non-significant.

Next, we tested if the P+T treatment is effective in ameliorating W024 and W121 SC-islet dysfunction (Figure 4E). WFS1 protein expression was restored in the treated SC-islets, as observed in both the W024 and W121 iPSCs (Figure 4F). In addition, the P+T treatment greatly inhibited cell death in the W024 and W121 SC-islets (Figure 4G). As expected with greater WFS1 protein, the W024 and W121 SC-islet functional capacity improved with P+T treatment, measured by insulin secretion in low and high glucose (Figure 4H). In summary, P+T treatment restored WFS1 expression and increased insulin secretion capabilities of W024 and W121 SC-islets.

### Cellular stress is mitigated by a combination treatment of 4-PBA and TUDCA in SC-islets with *WFS1* c.1672C>T, p.R558C variant

We performed multiplexed single-cell RNA sequencing (sc-RNA seq) using the 10x Genomics platform to investigate genotype-phenotype correlations and an efficacy of the P+T combination treatment on SC-*β* cells more precisely. We utilized Cell Hashing, which applied oligo-tagged antibodies to the cell surface proteins of individual samples, thus allowing for detection of individual samples within a pooled cell population (48). We sequenced four biological replicates per cell line, treated with or without P+T for 7 days. In total, we sequenced 16 samples with 8 samples in each pooled population which was submitted separately based on the cell line. In total, we sequenced 13,951 Stage 6 SC-islet cells differentiated from W024 and W121 iPSCs to study the effects of P+T treatment (W024: 2,619 cells, W024, P+T: 3,158 cells, W121: 3,625 cells, and W121, P+T: 4,549 cells, 4 biological replicates for each sample). The sc-RNA seq data was analyzed using dimensionality reduction and unsupervised clustering to classify individual cells into cell populations based on similarities in their transcriptome profiles. The cell types were identified by aligning the top upregulated genes in each cell cluster population with published pancreatic transcriptome data (49, 50). After identifying the *β* cell population in each sample, we combined the 2,329 SC-*β* cells from the four experimental conditions and performed Principal Component Analysis (PCA) and unsupervised clustering. The *β* cells clustered together based on genetic background, regardless of combination treatment (Figure 5A), suggesting the *β* cell transcriptional profile was not greatly changed in response to P+T.

**Figure 5.**
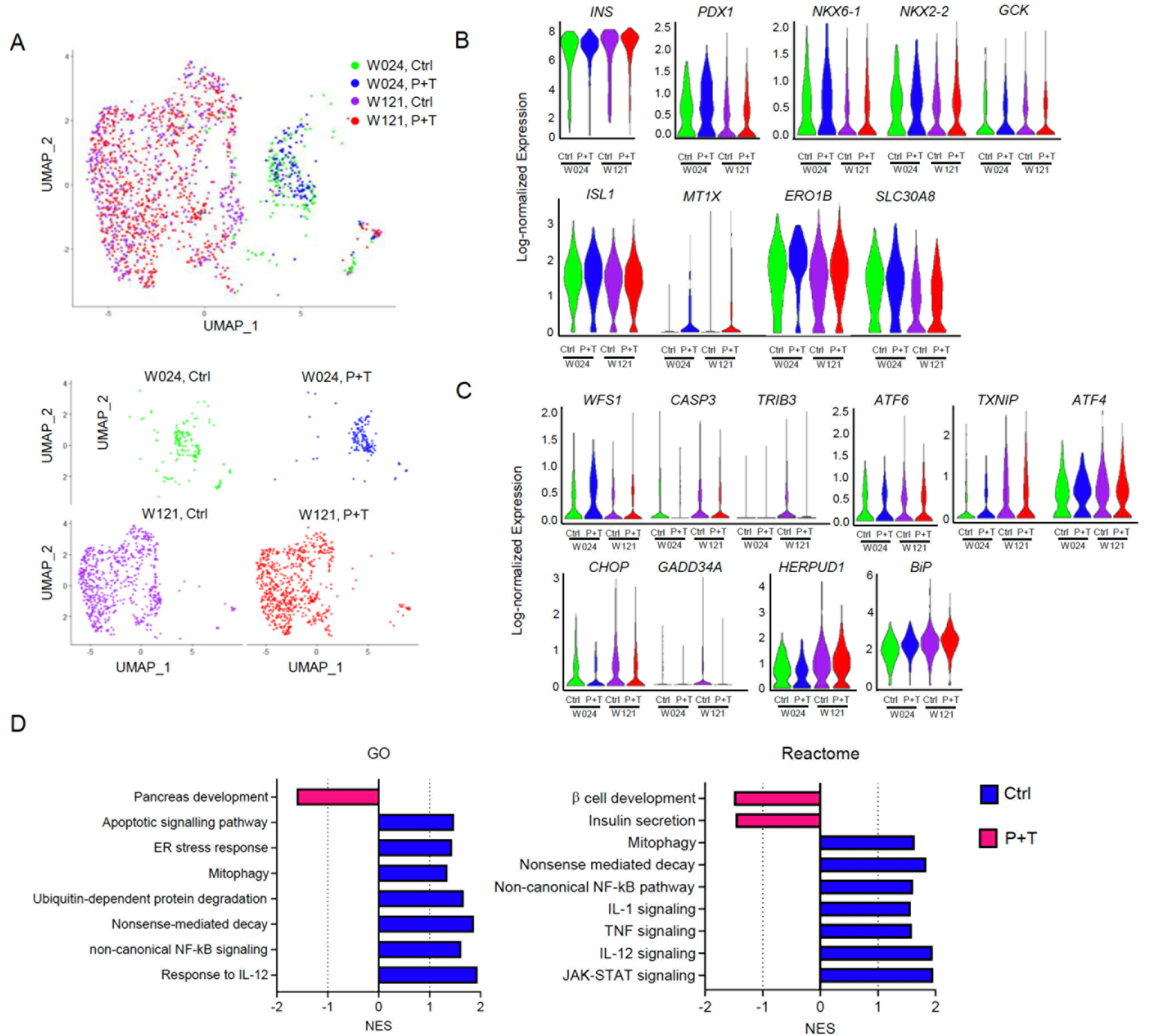
Single-cell transcriptional evaluation of a combination treatment with 4-PBA and TUDCA on SC-β cells. (A) Uniform Manifold Approximation and Projection (UMAP) plot from unsupervised clustering of combined transcriptional data from scRNA-seq of W024, Ctrl (green); W024, P+T (blue); W121, Ctrl (purple) and W121, P+T (red) SC-*β* cell populations. Lower plots are UMAP plots split by experimental conditions. (B) Violin plots detailing log-normalized gene expression of *β* cell genes in the same populations as (A). Log fold change and P values for violin plots are available in Supplemental Table S2. (C) Violin plots detailing log-normalized gene expression of ER stress and apoptotic genes in the same populations as (A). Log fold change and P values for violin plots are available in Supplemental Table S3. (D) GO and Reactome GSEA, quantified by the normalized enrichment score (NES), for pathways upregulated in the combined population of W024 and W121 SC-*β* cells treated with (red) or without (blue) P+T. NES values, P values, FDR q-values, and gene set lists are available in Supplemental Table S4.

Next, we evaluated the key *β* cell genes and ER stress markers. WFS1 pathogenic variants cause ER stress, resulting in altered expression of b cell genes in SC-*β* cells (51). Expression of the insulin gene (*INS*), crucial transcription factors for b cell differentiation (*ISL1*, *NKX6-1*, *NKX2-2* and *PDX1*), *β* cell maturation genes (*MT1X* and *ERO1B*), and a *β* cell function gene (*GCK*) were similar between untreated W024 and W121 SC-*β* cells (Figure 5B, and Supplemental Table S2). Interestingly, the expression of *SLC30A8*, an alternate *β* cell maturation gene, was significantly higher in W024 SC-*β* cells compared to W121 (Figure 5B, and Supplemental Table S2). The P+T treatment did not change the gene expression for many genes in W024 SC-*β* cells (Figure 5B, and Supplemental Table S2). On the other hand, *MT1X* and *ERO1B* were highly expressed in W121 SC-*β* cells treated with P+T compared to untreated (Figure 5B, and Supplemental Table S2). Unlike WFS1 protein levels in SC-islets, *WFS1* transcription within the SC-*β* cell population was similar between both W024 and W121 SC-*β* cells (Figure 4C, Figure 5C, and Supplemental Table S3). Some ER stress markers (*TXNIP* and *BiP*) were highly expressed in W121 SC-*β* cells compared to W024 (Figure 5C and Supplemental Table S3), whereas other ER stress markers (*ATF6*, *ATF4*, *CHOP*, *GADD34A*, *TRIB3* and *HERPUD1*) and apoptotic (*CASP*) genes were not statistical different between both W024 and W121 SC-*β* cells (Figure 5C and Supplemental Table S3). The expression of *WFS1*, ER stress markers, and apoptotic genes were not statistically altered by the P+T treatment in either W024 and W121 SC-*β* cells (Figure 5C, and Supplemental Table S3).

We performed gene set enrichment analysis (GSEA) on the SC-*β* cells. Gene sets pertaining to NMD, ubiquitination-mediated protein degradation, and oxidative stress were enriched in the untreated SC-*β* cells compared to the P+T-treated SC-*β* cells (Figure 5D, and Supplemental Table S4). Interestingly, we found the inflammation and the selective mitochondrial autophagy (mitophagy) pathways were also enriched in the untreated SC-*β* cell population (Figure 5D, Supplemental Figure 5, and Supplemental Table S4). Although we did not observe major changes in the expression of ER stress markers and apoptotic genes in W024 and W121 SC-*β* cells treated with P+T, gene sets pertaining to apoptosis and ER stress were enriched in the untreated SC-*β* cells (Figure 5D, Supplemental Figure 5, and Supplemental Table S4). Of note, gene sets related to insulin secretion, and *β* cell development were enriched in the P+T-treated SC-*β* cell population (Figure 5D, Supplemental Figure 5, and Supplemental Table S4). Additionally, gene sets related to regulation of cytosolic K^+^ and Ca^2+^ levels were increased, which play an important role in *β* cell differentiation and function (52–54) (Supplemental Figure 5, and Supplemental Table S4). Collectively, P+T treatment mitigated cellular stress increased by pathogenic *WFS1* variants without changing *β* cell identity, which resulted in increased *β* cell and insulin secretion in W024 and W121 SC-*β* cells.

### A combination treatment of 4-PBA and TUDCA delays the diabetic phenotype progression in *Wfs1* deficient mice

Finally, we verified the efficacy of our combination treatment with chemical chaperones with an *in vivo* study. The field lacks a c.1672C>T, p.R558C variant *WFS1* mutation mouse model. Therefore, we employed 129S6 whole body *Wfs1*-knockout (*Wfs1* KO) mice. This mouse model develops progressive glucose intolerance during adolescence, hence a mouse model of Wolfram syndrome (20). We confirmed *Wfs1* KO mice developed glucose intolerance at 5-6 weeks old (Supplemental Figure 6A). We treated the mice at 5-6 weeks old with food containing 4-PBA and TUDCA (4-PBA: 0.338% and TUDCA: 0.225%, refer to P+T chow) for one month. Both groups of *Wfs1* KO mice consumed similar amount of chow (Supplemental Figure 6B). After feeding for one month, *Wfs1* KO mice fed with control chow developed more severe glucose intolerance (Supplemental Figure 6C). Conversely, an intraperitoneal glucose tolerance test (IP-GTT) blood glucose curve was similar to the baseline outcome in *Wfs1* KO mice fed with P+T chow (Supplemental figure 6D), indicating that P+T chow delayed the progression of the diabetic phenotype. Collectively, we observed delays in the Wolfram diabetic phenotype in *Wfs1* KO mice when using the P+T treatment. Therefore, we expect the combination treatment to be efficacious against diabetic Wolfram phenotypes caused by the *WFS1* c.1672C>T, p.R558C variant *in vivo*.

## Discussion

In this study, we characterize a unique pathogenic *WFS1* c.1672C>T, p.R558C variant, which is associated with a mild form of Wolfram syndrome and highly prevalent in the Ashkenazi Jewish population. Molecular investigation revealed WFS1 p.R558C has a greater post-translational stability in cells compared to another pathogenic variant, *WFS1* c.2654C>T, p.P885L. Based on the molecular characterization of WFS1 p.R558C, we hypothesized a combination treatment of 2 chemical chaperones, 4-PBA and TUDCA, could provide a new treatment for Wolfram syndrome. We demonstrated that a combination treatment of 4-PBA and TUDCA increased WFS1 expression within the cell, ameliorating organelle dysfunction and the associated apoptosis. We also identified that patient SC-islets show genotype-phenotype relationships that correlate with clinical observations, and a combination treatment with 4-PBA and TUDCA improved insulin secretion in this cellular model of Wolfram syndrome.

Wolfram syndrome is a very rare genetic disorder. In the United Kingdom, the prevalence is 1:770,000 and the carrier frequency of pathogenic *WFS1* variants is 1:354 (4). We observed greater carrier frequency of the *WFS1* c.1672 C>T, p.R558C variant in the Ashkenazi-Jewish population with a frequency of 1:43 as compared to 1:76 in the Ashkenazi/Sephardi Jewish population and 1:2,268 in the Sephardi population Although present in the Sephardi population, as six carriers out of 13,608 Sephardi subjects tested, we suspect that the low frequency of *WFS1* c.1672 C>T, p.R558C variant might be explained by historical admixture with Ashkenazim. Instances of forgotten or suppressed admixture of Gentile and Jewish subjects has been previously shown in a study focusing on genome-wide Jewish genetic signature (55).

It was reported that WFS1 is subject to a ubiquitin-proteasomal degradation by Smurf1, a HECT-type E3 ubiquitin ligase, which recognizes the degron within the C-terminal luminal region of WFS1 (aa671-700) (56). Missense mutations in the degron or truncating mutations lacking the degron are resistant to Smurf1-mediated degradation (56). Other *WFS1* variants that retain the functional degron can lead to complete depletion or degradation of the WFS1 protein by NMD or proteasomal degradation (31, 32). Our study demonstrated WFS1 p.R558C protein is also subjected to proteasomal degradation. However, the degradation rate was lower than the WFS1 p.P885L variant. Additionally, WFS1 p.R558C protein was detected while conducting *in vivo* and *in vitro* experiments, indicating that a portion of the WFS1 p.R558C protein escapes proteasomal degradation. The remaining WFS1 p.R558C protein may compensate for the loss of normal WFS1 function. This hypothesis is consistent with the observation that clinically, patients carrying the homozygous *WFS1* c.1672 C>T, p.R558C variant have mild or less severe phenotypes of Wolfram syndrome. Our findings strongly suggest that alternate degron-retaining pathogenic *WFS1* variants require further protein expression characterization to determine the genotype-phenotype correlation.

Most of the studies conducting experimental genotype-phenotype correlation analysis employ overexpression of mutant WFS1 proteins in transfected cell lines such as HEK293T, HeLa, and COS7 (56, 57). This system is incapable of evaluating detrimental effects of pathogenic *WFS1* variants on physiological functions in disease-relevant cells. In pursuit of more reliable models and achieving a deeper understanding of Wolfram phenotypes, we have studied the patient derived disease-relevant cells. SC-islets, containing SC-*β* cells, differentiated from patient iPSCs with homozygous *WFS1* c.1672C>T, p.R558C variant (W024) displayed insufficient b cell function and similar differentiation efficiency compared to control SC-islets. W024 SC-islet phenotypes were milder than W121 SC-islets differentiated from the heterozygous *WFS1* c.1672C>T, c.2654C>T variant iPS cell line, indicating that our SC-islets recapture Wolfram phenotypes based on pathogenic *WFS1* variants and could be a great *in vitro* model for further genotype-phenotype correlation analysis and drug discovery. The diabetes stem cell field has recently leveraged this technology to correct the WFS1 pathogenic variant using CRISPR/Cas9 gene-edited SC-islets from a patient with Wolfram syndrome to reverse pre-existing diabetes in a mouse (51).

We demonstrate that a combination treatment of 4-PBA and TUDCA is efficacious against the *WFS1* c.1672C>T variant. Numerous studies have demonstrated that 4-PBA and TUDCA are promising for the treatment of diabetes mellitus and other ER stress-related neurodegenerative diseases (58–61). However, most chaperone treatment studies used an individual chaperone and a single dose treatment. A combination treatment with these compounds has rarely been attempted. Combining several compounds is a common cancer treatment strategy, aiming to yield additive or synergistical efficacy (62). Recently, a study reported that a combination treatment of 4-PBA and TUDCA resulted in slower functional decline of patients with Amyotrophic Lateral Sclerosis (ALS), an ER stress-related disease (63). In our study, 4-PBA and TUDCA increased not only WFS1 protein but also mRNA levels in the patient-derived iPSCs. We predict this increase of *WFS1* mRNA level is induced by 4-PBA activity as an HDAC inhibitor which selectively promotes gene transcription (41, 64). Alternatively, *WFS1* and ER stress marker expression in SC-b cells were not statistically altered by the combination treatment with 4-PBA and TUDCA in our scRNA-seq analysis. This could be due to much variability of the scRNA-seq data set. Genes not in a consistent steady-state manner causes variable detection which is commonly observed in ER stress-inducible genes (65, 66). Additionally, current scRNA-seq technology only detects approximately 10% of the cellular mRNA molecules, resulting in difficulties to detect genes with minute expression within a single cell (67). Therefore, the scRNA-seq data set may be limited in the number of gene transcript copies detected for ER stress makers and *WFS1*, thus inhibiting underestimating the impact of 4-PBA and TUDCA on ER stress in the SC-*β* cells.

GSEA revealed a combination treatment of 4-PBA and TUDCA downregulated mitophagy, ER stress, and apoptosis. Mitophagy mediates clearance of damaged mitochondria in the cells (68). Pathogenic *WFS1* variants have been reported to cause mitochondrial dysfunction (38, 39), and we confirmed a combination treatment of 4-PBA and TUDCA restored mitochondrial function in NPCs, implying that mitophagy could be reduced by preventing mitochondrial dysfunction. Additionally, several inflammatory pathways were reduced by a combination treatment of 4-PBA and TUDCA in GSEA. We and others recently reported elevated expression and serum levels of inflammatory cytokines in patients with Wolfram syndrome (27, 69). ER stress and mitochondrial dysfunction have been well known to contribute inflammation (70–73). It is possible that the several inflammatory pathways were downregulated as a consequence of reduced ER stress and restored organelle function by a combination treatment of 4-PBA and TUDCA. In summary, a combination treatment of 4-PBA and TUDCA mitigated various cellular stresses caused by *WFS1* c.1672C>T, p.R558C variant, resulting in improved insulin secretion.

We previously reported on the restoration of Wolfram phenotypes by correcting pathogenic *WFS1* variants with CRISPR/Cas9 in SC-islets derived from patients with typical Wolfram syndrome (51). When comparing the functional assessments and scRNA-seq analyses, a combination treatment of 4-PBA and TUDCA had a milder effect compared to CRISPR/Cas9 gene correction. This is expected because CRISPR/Cas9 correction directly intervenes on the cause of Wolfram syndrome pathogenesis. However, the combination treatment of 4-PBA and TUDCA as a therapeutic is advantageous for ease of administration. Furthermore, prolonged 4-PBA and TUDCA administration could delay additional symptom onset, including hearing loss and neurodegeneration, which typically manifests at later stages of the disease. Therefore, verifying efficacy of the combination treatment on other relevant cell types and pathogenic *WFS1* variants is warranted.

This study harnesses multiple iPSC-derived *in vitro* disease models and demonstrates efficacy of a combination treatment of 4-PBA and TUDCA. The benefits of this treatment may be applicable to the other genetic ER stress-related diseases such as Wolcott-Rallison syndrome. Moreover, our differentiation technology of *in vitro* disease models could be leveraged as a drug screening tool to discover new therapies or pave the way to personalized medicine strategies for patients with Wolfram syndrome.

## Materials and Methods

### Human Study Approval

Subjects, and their parents or legal guardians, as appropriate, provided written, informed consent before participating in this study, which was approved by the Human Research Protection Office at Washington University School of Medicine in St. Louis, MO (IRB ID #201107067).

### Ashkenazi Jewish samples and Genotyping

All participants provided written consent to be used for research purposes. The consent form included that patient material would be used for clinical testing and that excess material would be de-identified for use of research purposes to characterize single gene disorders in the Ashkenazi Jewish population. We genotyped 87,093 subjects of Jewish descent for the presence/absence of the c.1672C>T p.R558C variant in the *WFS1* gene. These samples were anonymously obtained through the Dor Yeshorim screening program from 2017 to 2021. This includes samples obtained from Dor Yeshorim screening locations around the world. Subjects who use the Dor Yeshorim screening program typically represent all levels of Jewish Orthodoxy (74). When subjects submit samples for carrier screening, written consent is obtained for excess sample to be anonymously used for research to better characterize single gene disorders in the Jewish populations. Detailed methods of carrier screening at the time of testing have been previously described (75).

### Animal Study Approval

Animal experiment was performed according to procedures approved by the Institutional Animal Care and Use Committee at the Washington University School of Medicine.

### Induced pluripotent stem cell (iPSC) line

To generate iPSCs, we obtained PBMCs from patients with Wolfram syndrome. The iPSC lines were generated by the Genome Engineering and iPSC Center (GEiC) at Washington University in St. Louis with Sendai viral reprogramming. Control iPSC lines BJFF.6 and AN1.1 were obtained from GEiC at Washington University in St. Louis.

### Chemical chaperones

We obtained 4-PBA and TUDCA from Amylyx Pharmaceuticals Inc. 4-PBA and TUDCA were dissolved in PBS and used at final concentrations of 500 mM and 50 mM, respectively. Control conditions were treated with only PBS. The treatment time is described in the figure legends. Stock reagents were kept at 4°C and made fresh every 2 weeks. For animal experiments, we ordered Envigo custom-made chow containing 4-PBA and TUDCA (4-PBA; 34 mg/kg and TUDCA; 23 mg/kg in Teklad global 18% protein rodent diet). The animals ate ∼4 g of diet per day, with a goal of 6 g 4-PBA / kg of body weight / day and 4 g TUDCA / kg of body weight/day. The same base diet without chemical chaperones was used as a control chow.

### WFS1-HiBiT cloning, thermal stability, expression assay and high-throughput screen

The coding sequences for WT, p.p885L, and p.R558C WFS1 in pDONR221 were a kind gift from Richard Suderman and David Chao. The coding sequences were subcloned into a pcDNA3.1(+) backbone downstream of the Gly-Ser-HiBiT-Gly-Ser tag, using InFusion PCR (Takara) as previously described (76). For the NanoLuc control vector, the coding sequence was PCR amplified from pFN31K (Promega) using forward: 5’-ACCCAAGCTGGCTAGCcaccATGGTCTTCACACTCGAAGATTTCG-3’ and reverse: 5’-GATATCTGCAGAATTCttattccatggcgatcgcgc-3’. The amplicon was gel purified and subcloned into the NheI and EcoRI sites of pcDNA3.1(+) using InFusion PCR. Thermal stability experiments were performed as previously described (33). In brief, 1×10^6^ HEK293T were transfected in a 6-well dish using 3 μg of plasmid and 6.25 mL of Lipofectamine 2000 per well. 48 h after transfection, cells were lifted and resuspended in CETSA buffer containing DPBS (with CaCl2 and MgCl2) plus 1 g/L glucose, 1X Halt protease inhibitor cocktail. Samples were aliquoted to PCR strips at 30 µL per tube then heated for 3.5 min using a pre-heated thermal cycler, allowed to equilibrate to room temperature, and 6 µL of 6% NP40 was added to each tube. Samples were incubated at room temperature for 30 min. Substrate was added to a final concentration of 100 nM 11S fragment and 0.5X furimazine (Promega, NanoGlo Substrate). Luminescence was measured on a ViewLux HTS reader (PerkinElmer) equipped with clear filters. For temperature rescue experiments, 2.5×10^5^ HEK293T cells were transfected with plasmids a 24-well plates, using Lipofectamine 2000 and OptiMEM according to the manufacturer’s instructions. Cells were returned to a 37 °C incubator (5% CO2, 95% RH) for 48 h. Then, plates were left at 37 °C or shifted to 30 °C for 24 h. All medium was removed from the wells and lysis buffer (PBS + 1% NP40 + 1x protease inhibitor cocktail) was added to each well and plate was rotated at room temperature for 30 min. The lysate was pipetted to mix and 10 μL was transferred to a low volume white walled 384-well plate. 5 μL of substrate was added to a final concentration of 100 nM 11S fragment and 0.5X furimazine (Promega, NanoGlo Substrate). Luminescence was measured on a ViewLux HTS reader (PerkinElmer) equipped with clear filters. For proteasome inhibition experiments, cells were treated with 1600 nM bortezomib for 24h before measuring WFS1-HiBiT levels as described above. For PBA and TUDCA experiments, HEK293T cells were transfected in a T25 flask using 7.5 μg of DNA and 15 μL Lipofectamine 2000. 48 h after transfection, cells were re-seeded to white-walled 384-well plates (Corning) at 1×10^4^ cells per well in 40 μL volume. After overnight incubation, cells were treated with PBA and TUDCA (prepared in PBS) or vehicle control (PBS). Samples were lysed by removing all medium from wells, adding lysis buffer (PBS + 1% NP40 + 1x protease inhibitor cocktail) and rotating the plate for 30 min at room temperature. Reporter protein levels were assessed by adding 11S and furimazine to 100 nM and 0.5x, respectively. Luminescence was measured using a ViewLux HTS reader. For the high-throughput screening of the NCATS Pharmaceutical Collection, HEK293T cells were transfected with HiBiT tagged WT or R558C WFS1 (T225 flask, 62.5 μg DNA, 135 μL Lipofectamine 2000).

48 hours after transfection, cells were re-seeded to 1536-well white walled plates (Aurora, cyclic olefin polymer) at 4000 cells per well in a 5 μL volume. Cells were incubated overnight at 37 °C. Small molecules or DMSO vehicle control were transferred (30 nL, 10 mM stock, 60 mM) using an arrayed pin tool (Wako Automation) and cells were incubated for 24 h. 1 μL of 6% NP40 solution (final concentration 1%) was added to each well and plates were incubated at room temperature for 20 min. 1 μL of substrate (final concentrations: 100 nM 11S, 0.5x furimazine) were added and luminescence was measured using a ViewLux HTS reader equipped with clear filters.

### Cycloheximide chase assay and western blotting

Plasmid expressing HA-tagged human WFS1 was transfected into HeLa cells using TransIT-X2® Dynamic Delivery System (Mirus Bio; MIR 6000). After 24 hours, the cells were treated with 250 μg/ml Cycloheximide (SIGMA; C4859) then harvested at the time indicated in Figure 2D. Protein was extracted using M-PER™ Mammalian Protein Extraction Reagent (Thermo; 78501) in all experiments. An equivalent amount of total protein was loaded onto the SDS-polyacrylamide gel. Proteins were probed with primary and corresponding secondary antibodies. Antibody details can be found in Supplemental Table S5.

### Neural progenitor cell (NPC) differentiation

Undifferentiated stem cells were plated down on tissue culture plastic in mTeSR1 (StemCell Technologies; 05850) and cultured in a humidified 5% CO_2_ 37°C tissue culture incubator. NPC differentiation was performed as described previously (77, 78). Briefly, 4-6 x 10^4^ cells per well were seeded onto V-bottom 96 well plate in NPC differentiation media suppled with 10 µM Y27632 (Tocris; 1254) to generate embryonic bodies (EBs) (day 0). On day 4, EBs were plated on poly-L-ornithine solution (PLO) (20 ug/ml) and Laminin (1 mg/ml) coated 6 cm dish in NPC differentiation media. On day 12, neural rosettes were detached from the dish using STEMdiff™ Neural Rosette Selection Reagent (StemCell Technologies; 05832) and dissociated cells (NPCs) were plated on Matrigel and Laminin (5mg/ml) coated 6 cm dish. NPC differentiation media consists of Neurobasal-A (Life technologies; 10888022), 1xB27 supplement without vitamin A (Life technologies; 12587), 1% non-essential amino acid (Life technologies; 11140050), 0.1% 2-Mercaptoethanol (Life technologies; 21985-023), 1% PenStrep (Life technologies; 15140122), 1% Glutamax (Life technologies 35050061), 10mM SB-431542 (Tocris; 1614) and 100nM LDN-193189 (Tocris; 6053). Media was changed every other day in the whole differentiation process. NPCs were maintained in STEMdiff™ Neural Progenitor Medium (StemCell Technologies; 05833). Passage was performed with Accutase (SIGMA; A6964) and media was changed every day.

### Real-time qPCR

Total RNA was isolated using RNeasy Kits (Qiagen, 74106) and cDNA libraries were generated using high-capacity cDNA reverse transcription kits (Applied Biosystems, 4368814). Relative amounts of each transcript were calculated by the ΔΔCt method and normalized to human β-actin. Quantitative PCR was performed with the Applied Biosystems ViiA7 using PowerUp™ SYBR™ Green Master Mix (Applied Biosystems, A25741). Primers used for qPCR are listed in Supplemental Table S6.

### Measurement of apoptosis

10,000 NPCs or 20,000 stage 6 cells per well were seeded onto Corning® 96-well Flat Clear Bottom White Polystyrene TC-treated Microplates (Corning, 3610). After the cells were treated with 4-PBA and TUDCA, cell viability followed by caspase 3/7 activity were measured using CellTiter-Fluor™ Cell Viability Assay kit (Promega, G6080) and Caspase-Glo® 3/7 Assay System (Promega, G8090), respectively. Caspase 3/7 activity was normalized to cell viability.

### Mitochondrial respiration studies

NPCs were seeded on Seahorse cell culture plate (Agilent, 101085-004) overnight, then treated with 4-PBA and TUDCA for 48 hours. Media was replaced with DMEM (Agilent, 103575-100) supplemented with 2.5 mM glutamine, 17.5 mM glucose and 1 mM sodium pyruvate and the plate was placed in a non-CO_2_ incubator at 37°C for 1 hour. The cell culture plate was placed in a Seahorse XFe96 Analyzer. Sequential injections of 3 µM oligomycin, 0.25 µM carbonyl cyande-4-(trifluoromethoxy) phenylhydrazone (FCCP), and 1 µM rotenone and 2 µM antimycin A were placed in the analyzer injection ports. All compounds were from a Seahorse XF Cell Mito Stress Test Kit (Agilent, 103015-100). Four OCR measurements were recorded for baseline and following each injection. Cells were lysed with 1% Triton in TE buffer and the Quant-iT Picogreen dsDNA assay kit (Invitrogen; P7589) was used to normalize OCR measurements to DNA (ng) for each well.

### SC-islet differentiation

SC-islet differentiation was performed similar to as we have previously reported (79). Undifferentiated stem cells were plated down on tissue culture plastic in mTeSR1 (StemCell Technologies; 05850) and cultured in a humidified 5% CO_2_ 37°C tissue culture incubator. Stem cell passaging occurred every 3-4 days, with TrypLE (Life Technologies; 12-604-039) used for single cell dispersion and Vi-Cell XR (Beckman Coulter) for counting. Undifferentiated stem cells were seeded at 0.52 x 10^6^ cells/cm^2^ in mTeSR1+ 10 µM Y27632 (Abcam; ab120129) on Matrigel-coated (Corning; 356230) plates. On stage 6 day 7, cells were single cell dispersed with TrypLE and seeded in a 6 well plate at 5 x 10^6^ cells per well with 4 mL of stage 6 ESFM per well. Cells continued culturing on an orbi-shaker (Benchmark) set at 100 RPM in a humidified tissue culture incubator at 5% CO_2_ 37⁰C. Supplemental Table S7 contains subsequent feeding schedule, media formulations, and differentiation factors. Assays were carried out between stage 6 day 9 and 20.

### Glucose stimulated insulin secretion (GSIS)

GSIS was performed similar to as we previously reported (80). Cells were collected and washed with KRB buffer (128 mM NaCl, 5 mM KCl, 2.7 mM CaCl2 1.2 mM MgSO4, 1 mM Na2HPO4, 1.2 mM KH2PO4, 5 mM NaHCO3, 10 mM HEPES (Gibco; 15630-080), and 0.1% BSA in water)) before transfer to transwells in 2 mM glucose KRB solution for a 1 hour equilibration period. Cells were in a humidified incubator at 37°C 5% CO_2_ for all incubations. Samples were transferred to another 2 mM glucose KRB solution for 1 hour and the supernatant was collected. Lastly, samples were transferred to 20 mM glucose KRB solution for 1 hour and the supernatant was collected. After all incubations, transwells were moved to TrypLE for 30 minutes at 37°C for single cell dispersion and counted for normalization.

### Immunocytochemistry (ICC) and flow cytometry

For ICC of NPCs, cells were plated on eight-well chamber slides (Thermo; 177402) coated with Matrigel and laminin (5 µg/mL) and fixed in 4% Paraformaldehyde (PFA) for 20 minutes. After blocking with blocking buffer (10% donkey serum, 1% BSA and 0.1% TritonX-100 in PBS) for 1 hour, cells were incubated with primary antibodies diluted in antibody dilution buffer (1% donkey serum, 1% BSA, and 0.1% TritonX-100 in PBS) overnight at 4°C and then stained with corresponding secondary antibodies diluted in antibody dilution buffer for one hour at room temperature. Cells were imaged with a fluorescent microscope.

For flow cytometry, single cells were fixed with 4% PFA for 30 minutes at room temperature after single-cell dispersion with TrypLE. Cells were rinsed with PBS and underwent incubation in ICC solution (5% donkey serum and 0.1% TritonX in PBS) at room temperature for 45 minutes. After washing with ICC solution twice, cells were filtered before analysis on either the LSRII flow cytometer and BD LSR Fortessa X-20 (BD Biosciences). Dot plots and percentages for data analysis were generated with Flow Jo. Antibody details can be found in Supplemental Table S5.

### Single-cell RNA sequencing preparation

Single-cell RNA sequencing was performed similar to as we previously reported (81). Cells were single cell dispersed in TrypLE and stained with hashtags. There were four biological replicates per condition. The samples were pooled while keeping individual cell lines (W024, W121) in different samples and submitted to Washington University in St. Louis Genome Technology Access Center for library preparation and sequencing using the Chromium Single Cell 3’ Library and Gel Bead Kit v3 (10x Genomics). Cells were sequenced on the NovaSeq 5000 (Illumina) at 26x98bp. For the feature library, we used custom 8bp dual same TotalSeqA-HTO indexes (TruSeqIDTdual-D807, CAGATCAT-CAGATCAT) and IDT HTO cDNA PCR additives. Hashtag antibody list is in Supplemental Table S8.

### Single-cell RNA sequencing analysis

Cells with hashtags were analyzed and demultiplexed using hashtag oligos with Seurat v3.2.1. Cells with high mitochondria genes and a low number of genes mapped to the human genome were filtered out. Each sample was normalized with *NormalizeData* and *FindVariableFeatures* functions to remove outlier genes with a scaled z-score dispersion. The cells were separated into 4 groups for further analysis: W121, Ctrl; W024, Ctrl; W121, P+T; and W024, P+T. The cells were clustered on UMAP plots, where cells with similar gene expression are proximally located based on PCA with *FindNeighbors* and *FindClusters* functions. UMAP plots for each group were generated using a resolution of 0.3 and 30 dimensions to identify the separated clusters. The clusters represent different cell types which we defined based on differential gene expression (*FindAllMarkers* function) and aligning the top differentially expressed genes to the pancreatic, endocrine, exocrine, and off target cell types. To visualize the UMAP plots, we used the functions *RunUMAP* and *DimPlot*. After identifying the b cell population for each group, we found the differential expression between the b cell populations using *FindAllMarkers*. Data is represented in violin plots using *VlnPlot*.

### Gene set enrichment analysis

Genes between control and treated sample groups were analyzed for Reactome and GO pathway enrichment by GSEA (4.0.2). Gene sets NES > 1.0 or < -1.0 and NOM p-val < 0.05 were considered as significantly enriched gene sets.

### Animal study

129S6 whole body *Wfs1*-knockout mice were a kind gift from Dr. Sulev Kõks. All animal experiments were performed according to procedures approved by the Institutional Animal Care and Use Committee at the Washington University School of Medicine (20-0334). 5-6 weeks-old female mice were used for this study. Food consumption rate was determined by monitoring D diet mass (g) per week. Animal number for each group is indicated in figure legends. Intraperitoneal glucose tolerance test (IP-GTT) was performed according to standard procedures of the NIH-sponsored National Mouse Metabolic Phenotyping Centers (http://www.mmpc.org).

### Statistical analysis

Statistical analysis was performed by unpaired and paired t tests and one- and two-way ANOVA with Tukey’s or Dunnett’s tests. Statistical tests are specified in figure legends. P < 0.05 was considered statistically significant. Data are shown as means ± SEM unless otherwise noted.

## Supplementary Materials

Supplemental Figure 1. Classification of pathogenic WFS1 variants and karyotypes of iPS cell lines used in this study

Supplemental Figure 2. High throughput screening of the NCATS Pharmaceutical Collection (NPC) to identify small molecules that increase R558C expression.

Supplemental Figure 3. P+T treatment reduced caspase 3/7 activity in NPCs derived from patients with typical Wolfram syndrome.

Supplemental Figure 4. A schematic of 6 stage SC islets differentiation protocol mimicking embryonic development of pancreatic endocrine cells.

Supplemental Figure 5. Gene set enrichment analysis (GSEA) on the SC-*β* cells.

Supplemental Figure 6. in vivo verification of a combination treatment with chemical chaperones.

Supplemental Table S1. Carrier frequencies for WFS1 c.1672C>T, p.R558C related to Figure 1.

Supplemental Table S2. Log fold change values for key *β* cell genes in Fig 5B.

Supplemental Table S3. Log fold change values for *WFS1* and ER stress markers in Fig. 5C.

Supplemental Table S4. Gene Set Enrichment Analysis (GSEA) details.

Supplemental Table S5: Antibody list used in this study.

Supplemental Table S6: Primer list used in this study.

Supplemental Table S7: Media formulation, Differentiation factors and protocol for SC-islets differentiation.

Supplemental Table S8: Hashtag antibody list used in this study.

## Acknowledgements

This work was partly supported by the grants from the National Institutes of Health (NIH)/NIDDK (DK112921, DK020579), NIH/ National Center for Advancing Translational Sciences (NCATS) (TR002065, TR000448) and philanthropic supports from the Silberman Fund, the Ellie White Foundation for the Rare Genetic Disorders, the Snow Foundation, the Unravel Wolfram Syndrome Fund, the Stowe Fund, the Eye Hope Foundation, the Feiock Fund, Associazione Gentian - Sindrome di Wolfram Italia, Alianza de Familias Afectadas por el Sindrome Wolfram Spain, Wolfram syndrome UK, and Association Syndrome de Wolfram France to F. Urano. Research reported in this publication was also supported, in part” by the Washington University Institute of Clinical and Translational Sciences grant UL1TR002345 from the NIH/NCATS. The content is solely the responsibility of the authors and does not necessarily represent the official view of the NIH. We thank all the members of the Washington University Wolfram Syndrome Study and Research Clinic for their support (https://wolframsyndrome.wustl.edu) and all the participants in the Wolfram syndrome International Registry and Clinical Study, Research Clinic, and Clinical Trials for their time and efforts. We would also like to thank Chinyere Onwumere in Washington University School of Medicine for her support concerning *in vivo* experiments, Dr. Kesavan Meganathan in Washington University School of Medicine for helping us to establish NPC differentiation, Dr. David Chao and Dr. Richard Suderman in Stowers Institute for Medical Research for providing the plasmids, and Dr. Sarah Gladstone and Ms. Beth White for scientific feedback. WY, AS, and MJH were supported by the intramural research program of the NCATS, NIH.

## Author contributions

RAK, KGM, JRM, and FU conceived the experimental design. YH, MMJ, TW, JE, and FU conducted clinical investigation. WY, MJH and AS performed WFS1 SplitLuc assays and screening. KGM differentiated *β* cells and performed functional assays. KGM and PA performed sequencing analysis. RAK and CMB performed animal experiments. RAK differentiated NPC and RAK, CMB, and KK performed the other in vitro experiments. JC, JK, and KL examined data on 4-PBA+TUDCA. JRM and FU supervised the data. RAK, KGM, and FU wrote the manuscript. All authors edited and reviewed the manuscript.

## Data and materials availability

All data associated with this study are present in the paper or the Supplementary Materials. iPSC lines are available under a material transfer agreement upon request to the corresponding authors.

