## Supplemental Material for "Genotype-phenotype correlation analysis and therapeutic development using a patient stem cell-derived disease model of Wolfram syndrome"

Supplementary Materials

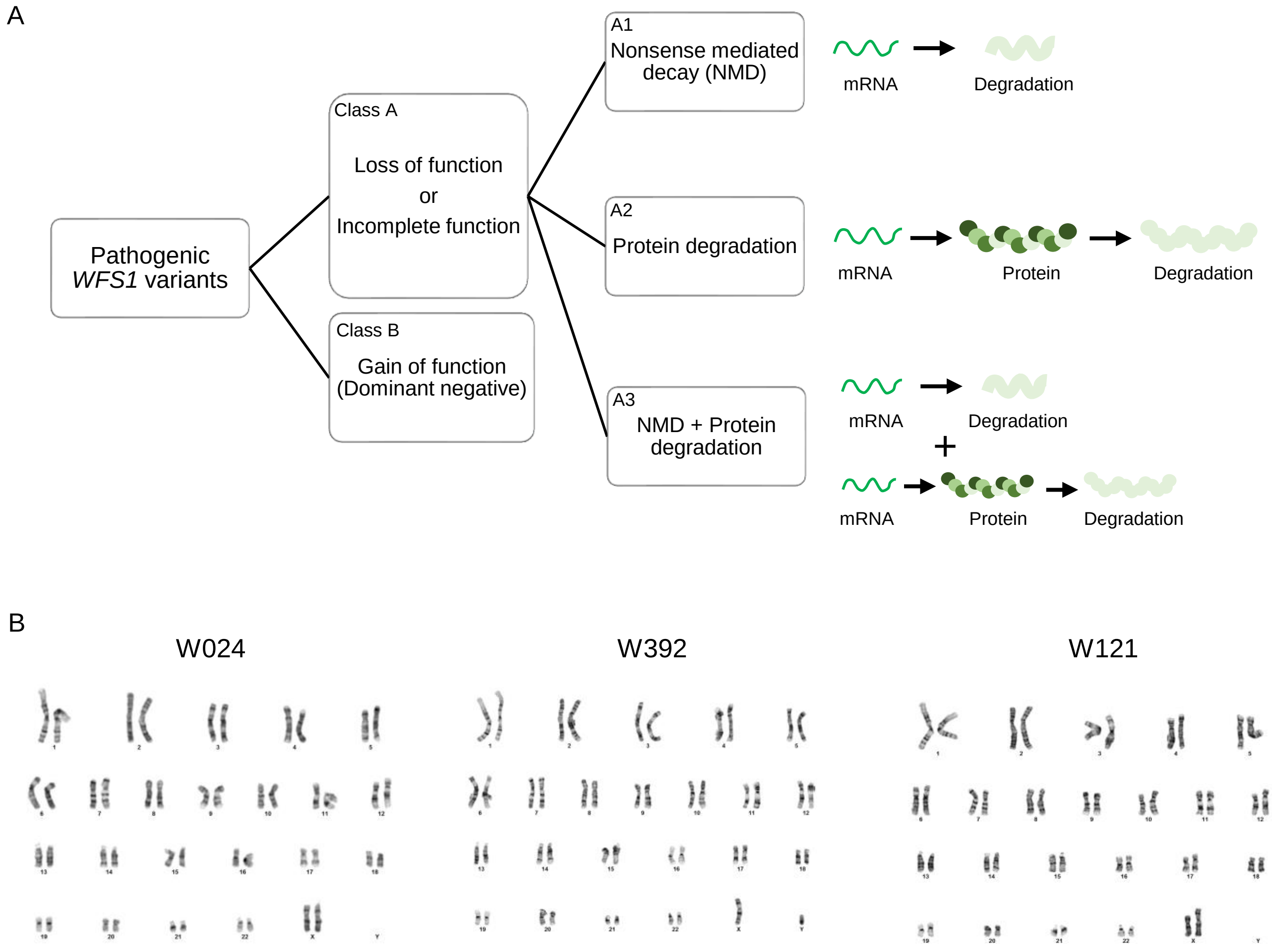


**Supplemental Figure 1. Classification of pathogenic WFS1 variants and karyotypes of iPS cell lines used in this study**

(A) A schematic of classification of pathogenic WFS1 variants in terms of protein expression. (B) Normal 46XX (W024 and W121) and 46XY (W392) karyotypes of derived iPS cell lines.

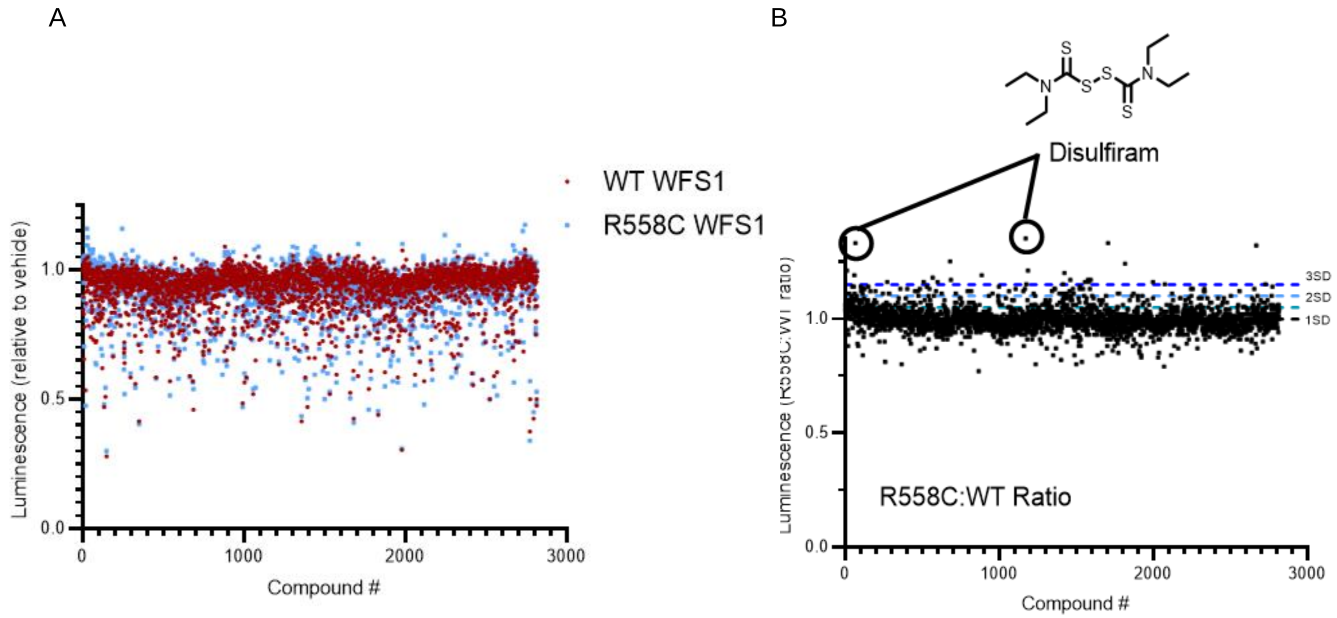


**Supplemental Figure 2. High-throughput screening of the NCATS Pharmaceutical Collection (NPC) to identify small molecules that increase R558C expression.**

(A) Expression of HiBiT tagged WFS1 variants after 24 h treatment with compounds (60 μM). The effect of each compound for WT (red dot) and R558C (blue dot) is shown. (B) R558C selectivity was assessed by calculating the ratiometric effect for R558C relative to WT-WFS1. Disulfiram, was the top hit (two samples contained in the library).

A

| Pt # | Age | Sex | Allele 1 *WFS1* | Allele 2 *WFS1* | Diabetes | Optic atrophy |
|---|---|---|---|---|---|---|
| W4 | 23.8 | Female | c.1112G>A; p.W371X | c.1885C>T; p.R629W | 2.3 | 5 |
| W9 | 14.3 | Male | c.376G>A; p.A126T | c.1838G>A; p.W613X | 10.8 | 11 |
| W13 | 5.9 | Female | c.599delT; p.L200fs286Stop | c.2254G>T; p.E752Stop | 4.8 | 5.2 |
| W15 | 10.8 | Female | c.439delC; p.R147fsX163 | c.1620G>A; p.W540X | 2.8 | 7 |

B

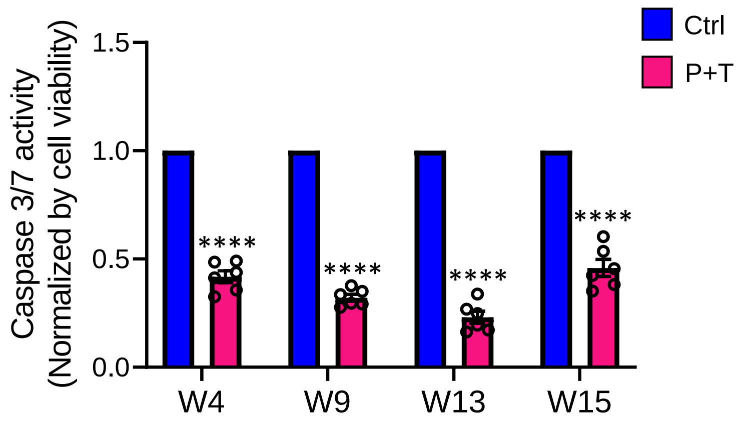


**Supplemental Figure 3. P+T treatment reduced caspase 3/7 activity in NPCs derived from patients with typical Wolfram syndrome**

(A) Information on the four patients with typical Wolfram syndrome, including the genetic location of autosomal recessive pathogenic variants in *WFS1* and the onset age of symptoms. (B) Caspase 3/7 activity normalized with cell viability in NPCs treated with or without P+T for 24 hours. (n=6, ****P < 0.0001 by unpaired t-test compared to Ctrl)

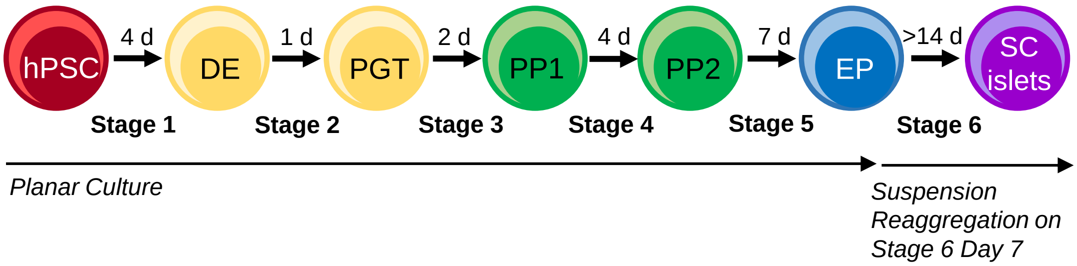


**Supplemental Figure 4. A schematic of 6 stage SC-islets differentiation protocol mimicking embryonic development of pancreatic endocrine cells**

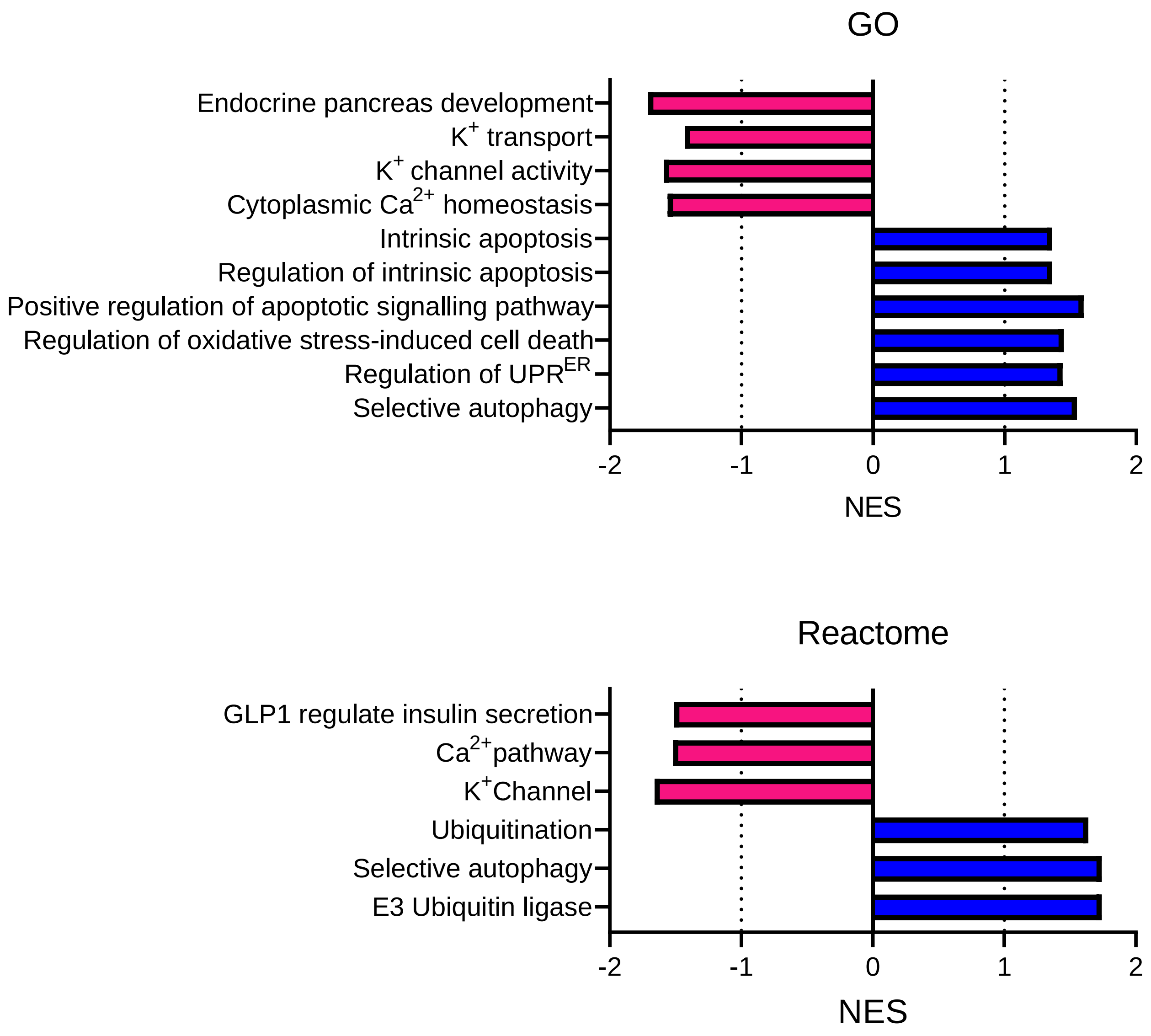


**Supplemental Figure 5. Gene set enrichment analysis (GSEA) on the SC-β cells**

GO and Reactome GSEA, quantified by the normalized enrichment score (NES), for pathways upregulated in the combined population of W024 and W121 SC-β cells treated with (red) or without (blue) P+T. NES values, P values, FDR q-values, and gene set lists are available in Supplemental Table S3.

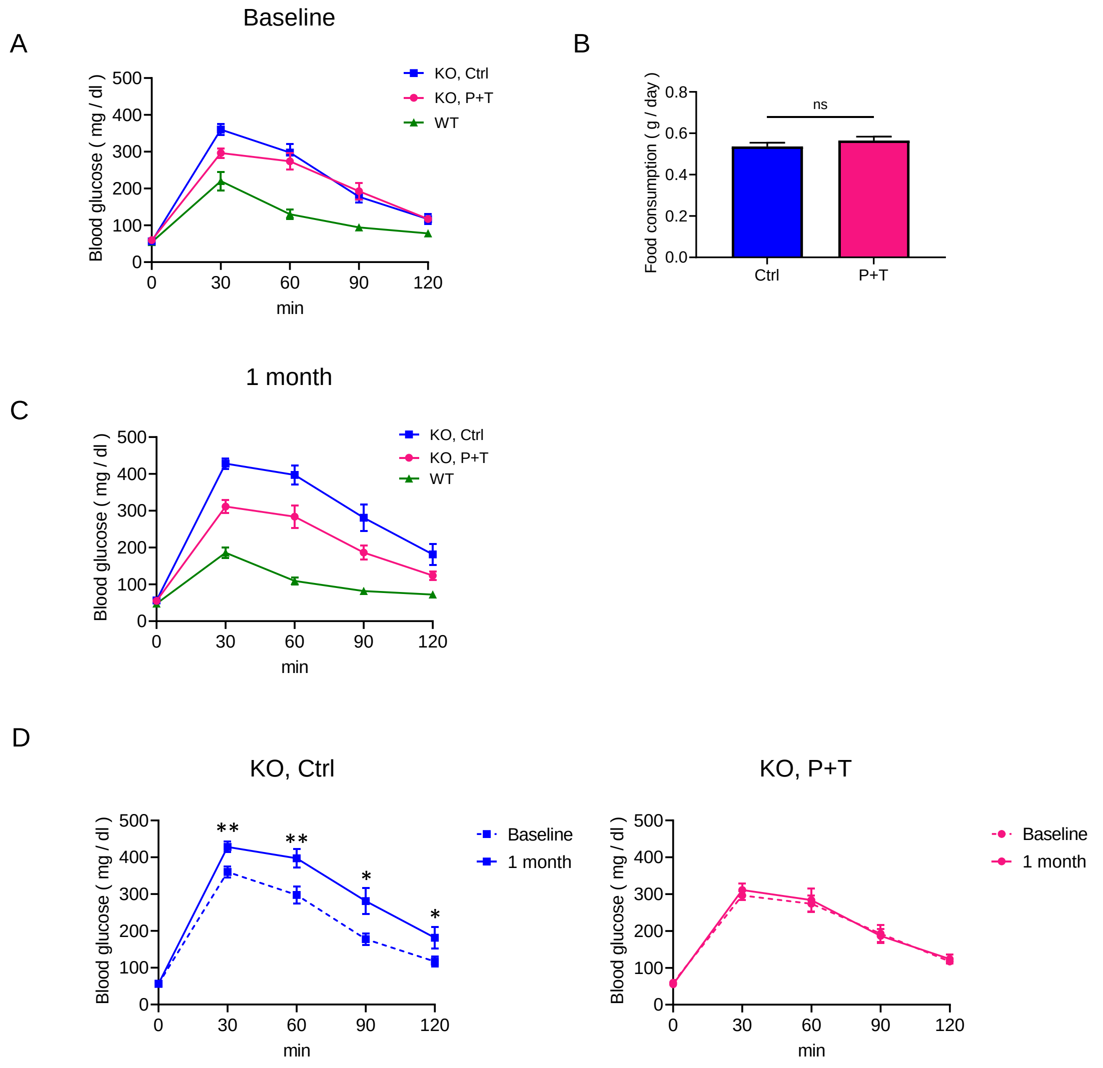


**Supplemental Figure 6. *in vivo* verification of a combination treatment with chemical chaperones**

(A) IP-GTT with WT or *Wfs1* KO mice at 5-6 weeks old before feeding with either Ctrl or P+T chow. (B) Food consumption rate in *Wfs1* KO mice fed with either Ctrl or P+T chow. (C) IP-GTT with WT or *Wfs1* KO mice fed with either Ctrl or P+T chow for 1 month. (D) IP-GTT with *Wfs1* KO mice fed with either Ctrl or P+T chow comparing between baseline and 1 month after feeding. KO Ctrl: n=11, KO P+T: n=12, *P < 0.05 and **P < 0.01 by unpaired t-test performed on each time point.
